# Mechanical transmission of Dengue Virus by *Aedes aegypti* may influence disease transmission dynamics during outbreaks

**DOI:** 10.1101/2023.03.07.531453

**Authors:** Hsing-Han Li, Matthew P. Su, Shih-Cheng Wu, Hsiao-Hui Tsou, Meng-Chun Chang, Yu-Chieh Cheng, Kuen-Nan Tsai, Hsin-Wei Wang, Guan-Hua Chen, Cheng-Kang Tang, Pei-Jung Chung, Wan-Tin Tsai, Li-Rung Huang, Yueh Andrew Yueh, Hsin-Wei Chen, Chao-Ying Pan, Omar S. Akbari, Hsiao-Han Chang, Guann-Yi Yu, John M. Marshall, Chun-Hong Chen

## Abstract

The escalating number of dengue virus (DENV) outbreaks and their worldwide spread pose a major threat to global public health. DENV transmission dynamics significantly influence outbreak duration and magnitude. Conventional DENV transmission requires an incubation period between mosquitoes biting infected humans and the mosquitoes becoming infectious. However, the possibility of immediate, mechanical transmission of DENV without viral replication in the mosquito has received little attention despite its potential importance.

Here, we show that *Aedes aegypti* mosquitoes can mechanically transmit DENV to susceptible mice immediately after biting infected mice without the need for an incubation period. By incorporating parameters from our experiments into a newly developed mathematical model, we found a significant impact on DENV outbreak characteristics.

Mechanical transmission may amplify existing disease transmission routes and influence outbreak dynamics. Our findings have implications for vector control strategies that target mosquito lifespan and suggest the possibility of similar mechanical transmission routes in other disease-carrying mosquitoes.

## Introduction

Global transmission of dengue virus (DENV) has become a significant public health concern over the past 40 years^1^. Almost half of the world’s population is now at risk of infection^2^, with over 390 million cases reported each year^3^. Recent years have seen large and rapid outbreaks of dengue occurring in multiple East Asian countries^4–6^, placing severe strain on local medical systems. This is in part due to the limited treatment options available, with DENV infection potentially causing a range of symptoms varying from mild flu-like features to severe dengue hemorrhagic fever and dengue shock syndrome^7^.

Given the paucity of treatment options, disease control heavily relies on interventions targeting the vectors themselves (*Aedes aegypti* [*Ae. aegypti*] and *Aedes albopictus*)^1^. However, DENV outbreaks can still occur within extremely short time periods with even relatively small infectious mosquito populations in dengue- receptive areas^8^. Although there is some evidence that an elevated density of mosquitoes plays a contributing role^5, 9^, a much-improved understanding of DENV transmission routes is necessary to identify methods to slow these outbreaks.

Multiple possible viral transmission routes exist, including air spray, fluid exchange, food, or insect vectors, and the relevant transmission route for each disease depends on specific pathogen features. These transmission routes greatly affect the infected population size during an epidemic. DENV transmission is assumed to follow a human-mosquito-human cycle^7^. Here, mosquitoes become infected via a blood meal from an infected human, with DENV replicating in the mosquito midgut for 3–5 days before needing up to 14 days to move to the salivary gland. The mature virus is then able to infect susceptible humans via a bite from this now-infected mosquito^7^.

This time period between the mosquito becoming infected and it becoming infectious, denoted as the extrinsic incubation period (EIP), has thus been estimated at approximately 7–14 days. The EIP influences the proportion of mosquitoes that become infectious after exposure to DENV, meaning it plays a key role in predicting the magnitude of dengue outbreaks. This property makes it useful for modeling dengue transmission dynamics^10, 11^, though parameter estimates for the EIP are variable^12^.

Regardless of the exact EIP value used in specific models, the assumption of a minimum necessary time period before mosquitoes become infectious is widespread; it is presumed that for time values below this EIP, DENV transmission is not possible. Thus, whilst these models are extremely useful, they are unable to account for alternative DENV transmission methods that do not require extrinsic incubation of the virus.

Interestingly, mechanical transmission (MT) of some viruses without an incubation period has been demonstrated in *Ae. aegypti*, including Chikungunya, poxviruses, myxoma virus, and lumpy skin disease virus (LSDV), between animals such as rabbits, sheep, and goats^13, 14^. DENV transmission might therefore also be transferred mechanically, as *Aedes* mosquitoes may require up to four bites (potentially from multiple humans) to take a complete blood meal^15, 16^, though it is unclear if/how MT contributes to dengue outbreaks.

To explore the potential importance of mechanical transmission (MT) as a route of DENV transmission, we conducted a mixture of simulations and laboratory experiments. Our simulations incorporating MT as a potential transmission route were similar to real-world data collected from Kaohsiung city. In our needle-sharing experiment, we found that needles puncturing DENV-infected mice could transmit the virus to naive mice. Furthermore, our sequential blood feeding assay demonstrated that *Ae. aegypti* mosquitoes feeding on DENV-infected mice were able to immediately transmit the virus to naive mice without an EIP. Our mathematical model incorporating parameters derived from these experiments found that MT could significantly impact the dynamics of DENV outbreaks.

Overall, our research highlights the importance of considering MT as a possible transmission route for DENV and its implications for vector control interventions. Our findings have important implications for worldwide mosquito control programs, as these programs typically focus on shortening the lifespan of *Aedes* mosquitoes, which may not be effective in preventing mechanical transmission. Our model can be used as a sentinel assay for evaluating the severity of DENV outbreaks.

## Results

### Mathematical models incorporating mechanical transmission suggest an influence on DENV transmission dynamics

The cycle of dengue transmission revolves around humans and mosquitoes. How mosquitoes transmit the virus from infected humans to healthy individuals is a critical aspect of DENV spread, given the multiple biting nature of *Aedes* blood feeding; MT may act to supplement biological transmission routes of infection, boosting the number of infected individuals in the early stages of outbreaks (Figure 1).

**Figure 1.**
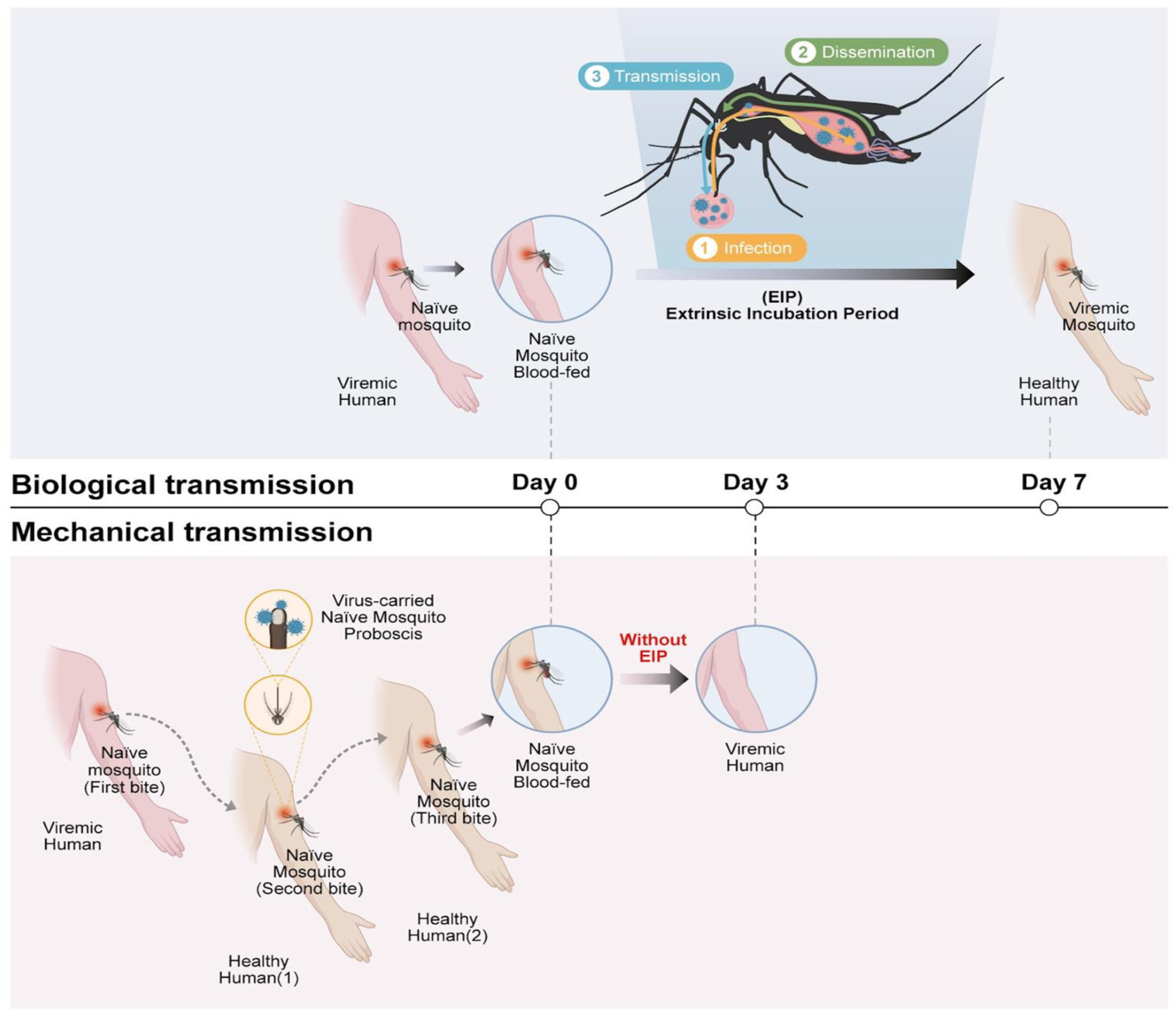
Mechanical transmission of DENV may alter disease outbreak dynamics and influence the time between infections. Diagrams of biological (upper panel) and mechanical (bottom panel) DENV transmission routes, with simulated data shown to the right demonstrating resulting changes in the speed and extent of disease spread.

The household is a significant focus for DENV transmission, and household level infection studies offer possible insight into the real-world impact of MT. Spatial and temporal studies indicate many dengue clusters occur in the same household with infection times of less than 7 days, far below the lowest possible EIP estimates. Indeed, DENV clusters (defined as two or more cases within 14 days within 150 m of each other) are highly focal in space and time^17, 18^ and have even been observed within the same household^19^. Though there are a number of factors which could contribute to this clustering of cases within time windows less than the minimum EIP estimate, MT may act as one component^14, 20^.

We first attempted to estimate MT’s importance by creating a mathematical model comparing either pure biological transmission, or a mixture of the two transmission types. Figures 2 A-D show the simulated daily incidence of DENV infections over the course of an epidemic for different probabilities of transmission per bite via MT, beginning with a single infected individual in one household of five individuals. The average duration of MT was assumed to be one hour, during which time a mosquito was expected to bite at least one other person (n_M_ = 1 - 4) with a probability of mechanical transmission per bite, p_M_, ranging from 100% to 10%. The mosquito biting rate ranged from 0.5 - 2, and the average number of adult female mosquitoes per person was set at 2. The total daily incidence with and without MT is partitioned into incidence due to standard transmission and incidence due to MT.

**Figure 2.**
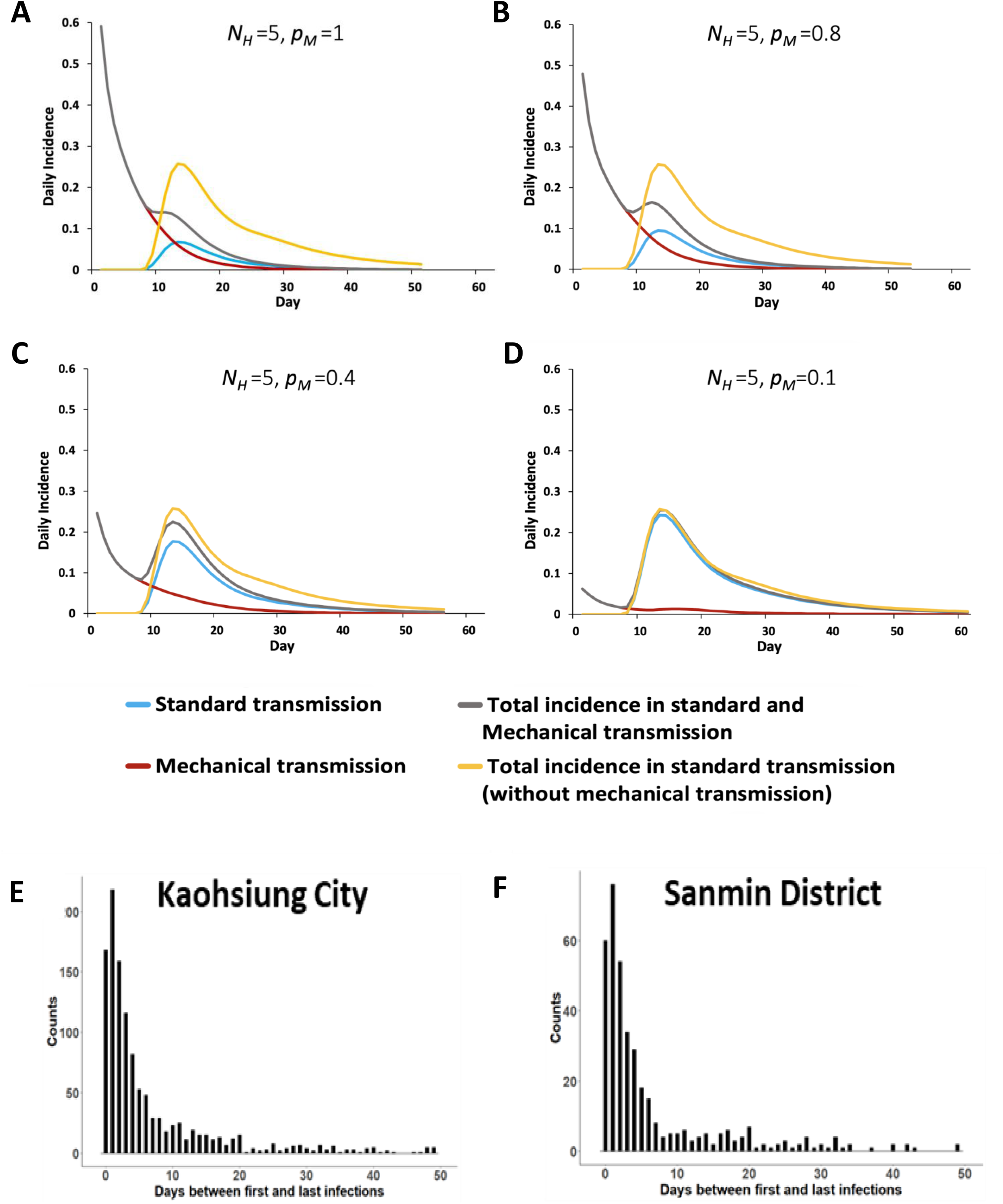
Mechanical transmission of DENV may alter disease outbreak dynamics and influence the time between infections. (A-D) Simulated daily incidence over the course of a dengue epidemic beginning with a single infected individual in an immunologically naïve population of 5 individuals (model in Equations 1–12 in Supplementary Figure 3 and parameters in Supplementary Table 1). The average duration of MT is one hour, during which time a mosquito is expected to bite at least one other person, nM = 1∼2, with a probability of mechanical transmission per bite, pM, of (A) 100%, (B) 80% (C) 40% or (D) 10%. The mosquito biting rate, b, was set to 0.5∼2, and the average number of adult female mosquitoes per person, k, was set to 2, 2.5, 3 or 5. The total daily incidence with MT (black line) or without MT (yellow line) is partitioned into incidence due to standard transmission following viral amplification in the vector (blue line) and incidence due to MT (red line). (E-F) Counts of cases throughout (E) all Kaohsiung City and (F) Sanmin district based on the number of days between first and last infections within a household during the 2015 Kaohsiung outbreak.

In all cases, provided p_M_ was greater than 0, cases of DENV occurred prior to the end of the necessary EIP. Figure 2C acts as an illustration of our simulation results, with the probability of a new individual becoming infected in the same household on day 2 being greater than 84%, assuming an 80% probability of MT per bite. Furthermore, at least one of the five individuals became infected within three days. These results suggest that MT can significantly impact the spread of dengue outbreaks, especially in households or other settings where mosquitoes have access to multiple susceptible hosts. Details of the model and parameter ranges used in our simulations can be found in the Methods section and Supplementary Tables 1 and 2.

To compare these simulations with real world data, we investigated DENV transmission dynamics using a detailed dataset of human infections recorded from Kaohsiung city, Taiwan during the 2015 outbreak. This outbreak resulted in tens of thousands of infections and affected the entire city, with a particularly severe impact in Sanmin district. Studying dengue clusters not only in the entire city but also specifically in this district, we found a staggering number of cases reported within a household less than 3 days after the initial reported infection (Figures 2E-F). Assuming a necessary EIP of at least several days, MT may be contributing towards this rapid spread of cases.

### Mechanical Transmission of DENV is possible in mice models

To investigate the potential of mechanical transmission (MT) of DENV, we conducted experiments using the cosmopolitan strain DENV-2 (TW2015) and immunocompromised AGB6 mice. These mice lack type I and type II interferon receptors^21^, enabling robust replication of the virus in their spleens, lungs, and intestines^12^. Utilising needle-stick assays, we tested whether DENV could be transferred via sharing of contaminated needles (Supplementary Figure 1A). Viremia, weight loss, and lethality occurred in mice that received the DENV-carrying needles, but not in those that received sterile needles, demonstrating the potential of MT of DENV (Supplementary Figure 1B-D).

We then moved on to investigate the possibility of transmission through the female proboscis of *Ae. aegypti* mosquitoes. We starved individual female mosquitoes for 24 hours and then allowed them to bite infected mice which had been injected with DENV three days prior. After partial completion of a blood meal, infected mice were replaced with naïve mice, which were bitten by female mosquitoes to complete their blood meals (Figure 3A).

**Figure 3.**
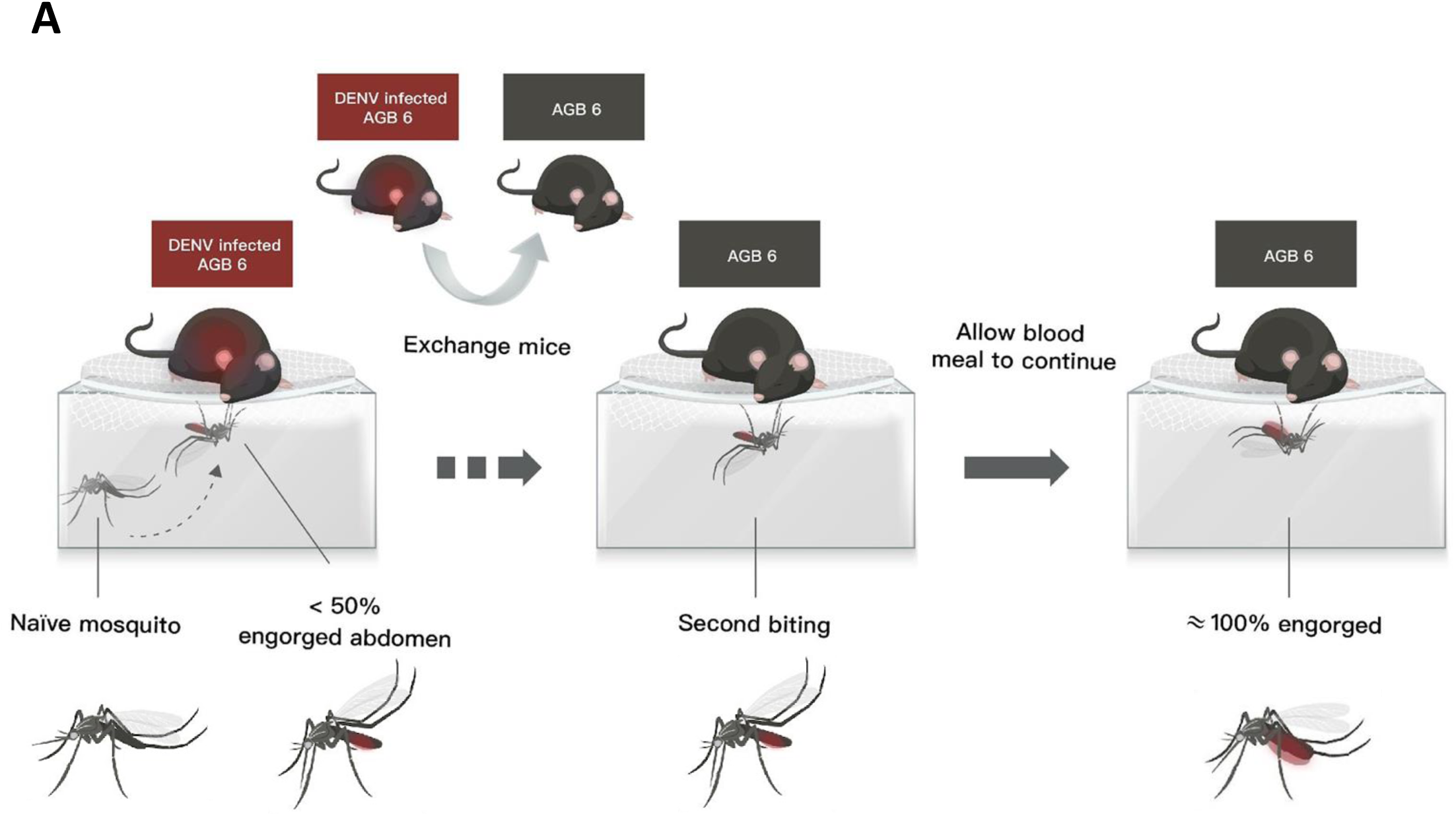
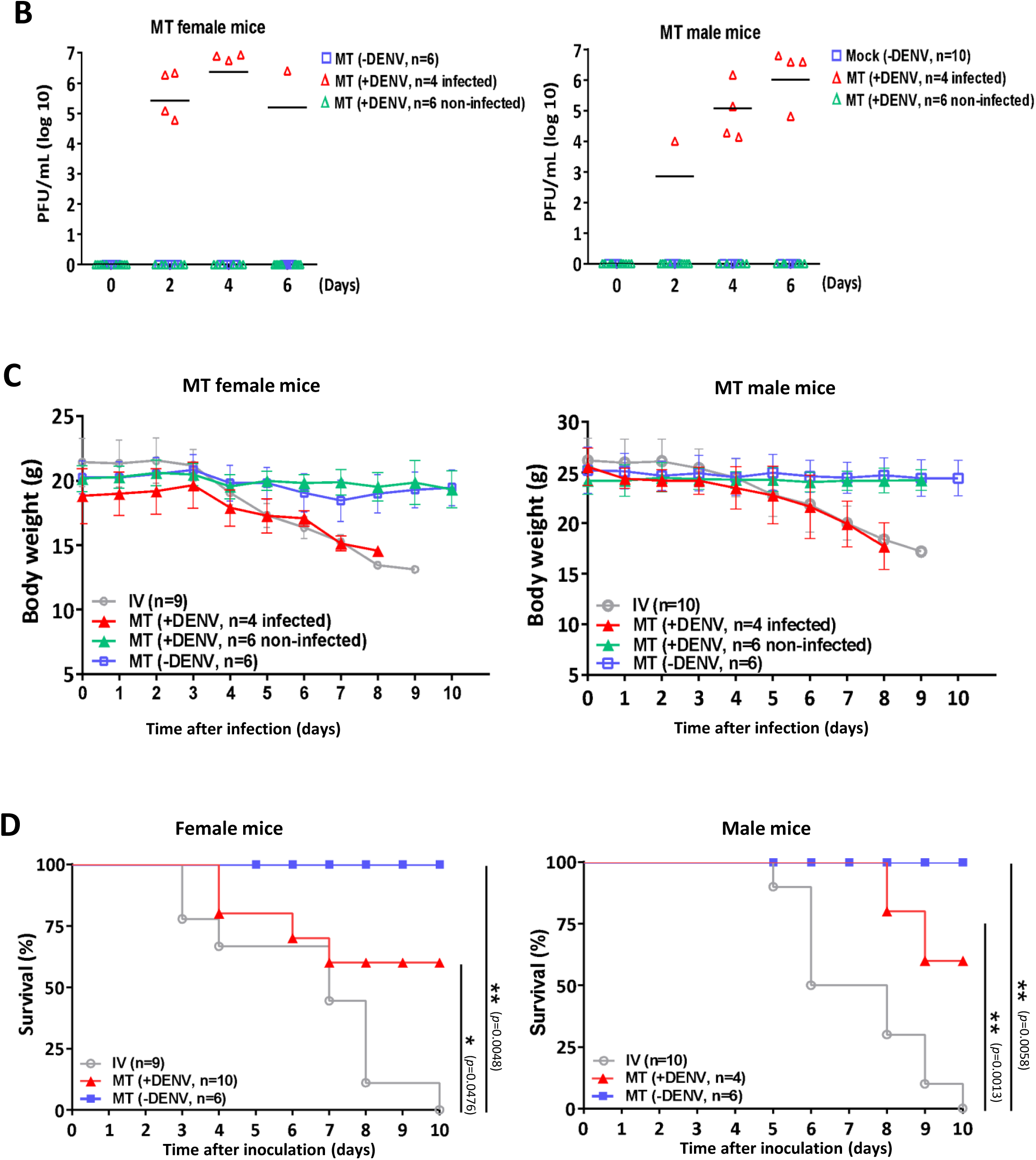
Effects of the mechanical transmission of DENV to mice by *A. aegypti*. (A) Schematic model to investigate mechanical transmission of DENV by *Aedes aegypti* mosquitoes. Mouse model in which *A. aegypti* mosquitoes at most 50% engorged with blood were used as a vector for DENV transmission via mechanical biting without biological DENV replication. Step one. AGB6 mice (red) were intravenously infected with DENV. Step two. Mosquitoes were allowed to feed on these mice until at most half-engorged, after which the mice were switched with naïve AGB6 mice (gray). Step three. The mosquitoes continued feeding, infecting the naïve AGB6 mice (referred to as MT mice) via mechanical transmission.(B-D) Serum DENV titer evaluated by plaque-formation assay (B), body weight (C) and survival (D) were examined in female (n=10) and male (n=10) mice infected with DENV or in mice (n=6 for male or female individually) without DENV infection via mechanical transmission (MT) by four bites of DENV-exposed *A. aegypti* or *A. aegypti* alone or in mice (n=9 for female or n=10 for male) infected with DENV with intravenous injection. Kaplan-Meier survival curves are shown in (D). Data is presented as mean±standard deviation for (C). Number of mice used in each group per individual experiment are highlighted. *p<0.05; **p<0.01. Details of the statistical analysis are provided in the Methods.

As for the needlestick experiment, naïve male and female mice bitten by infected mosquitoes could become infected (40% infectivity rate) as evidenced by detectable DENV serum titers (Figure 3B). Successful transmission of DENV was associated with a loss of body weight^12, 22^, as seen in mice that were either infected through MT or through direct tail vein inoculation (Figure 3C). Infected mice also died (Figure 3D), confirming the existence of MT as a lethal method of transmission for DENV. MT of DENV is thus possible by naïve mosquitoes immediately following incomplete biting of infected hosts, with this transmission sufficiently significant as to induce mortality.

### Residual DENV titers in the mosquito proboscis act as a determinant for MT success

Although HIV transmission through contaminated needles is well-known^23^, the minimum infectious dose remains unclear. Therefore, while we hypothesized that DENV in the mosquito proboscis could be responsible for infecting naïve mice, a sufficient concentration of the virus must be required for an infection to occur. This is supported by individual mosquito biting behavior observations, which showed that inadequate bites did not cause obvious symptoms of DENV infection (such as lethality, body weight loss, and serum DENV titer) in naïve AGB6 mice (Supplementary Figures 2A-D). Hence, we aimed to confirm the presence of DENV in the partially fed mosquito proboscis and investigate the minimum titers required for infection.

Initially, using RT-qPCR, we identified DENV genomic RNA in the proboscis of females after a single incomplete bite from DENV-infected mice (estimated 10^5^ copies) (Figure 4A). Subsequently, we aimed to correlate the detrimental effects resulting from MT with the residual DENV titer found within the mosquito proboscis. To achieve this, we dissected proboscis samples from *Ae. aegypti* that had obtained incomplete or complete blood meals from biting DENV-infected mice. These proboscis extracts were tail-vein injected into AGB6 naïve mice to test DENV virulence. Notably, we detected infectious DENV in the serum after intravenous injection (Figure 4B and 4E), which was associated with body weight loss (Figure 4C and 4F) and lethality (Figure 4D and 4G). Furthermore, the severity of these symptoms was closely correlated with viral proboscis extract viral loads in a dose-dependent manner.

**Figure 4.**
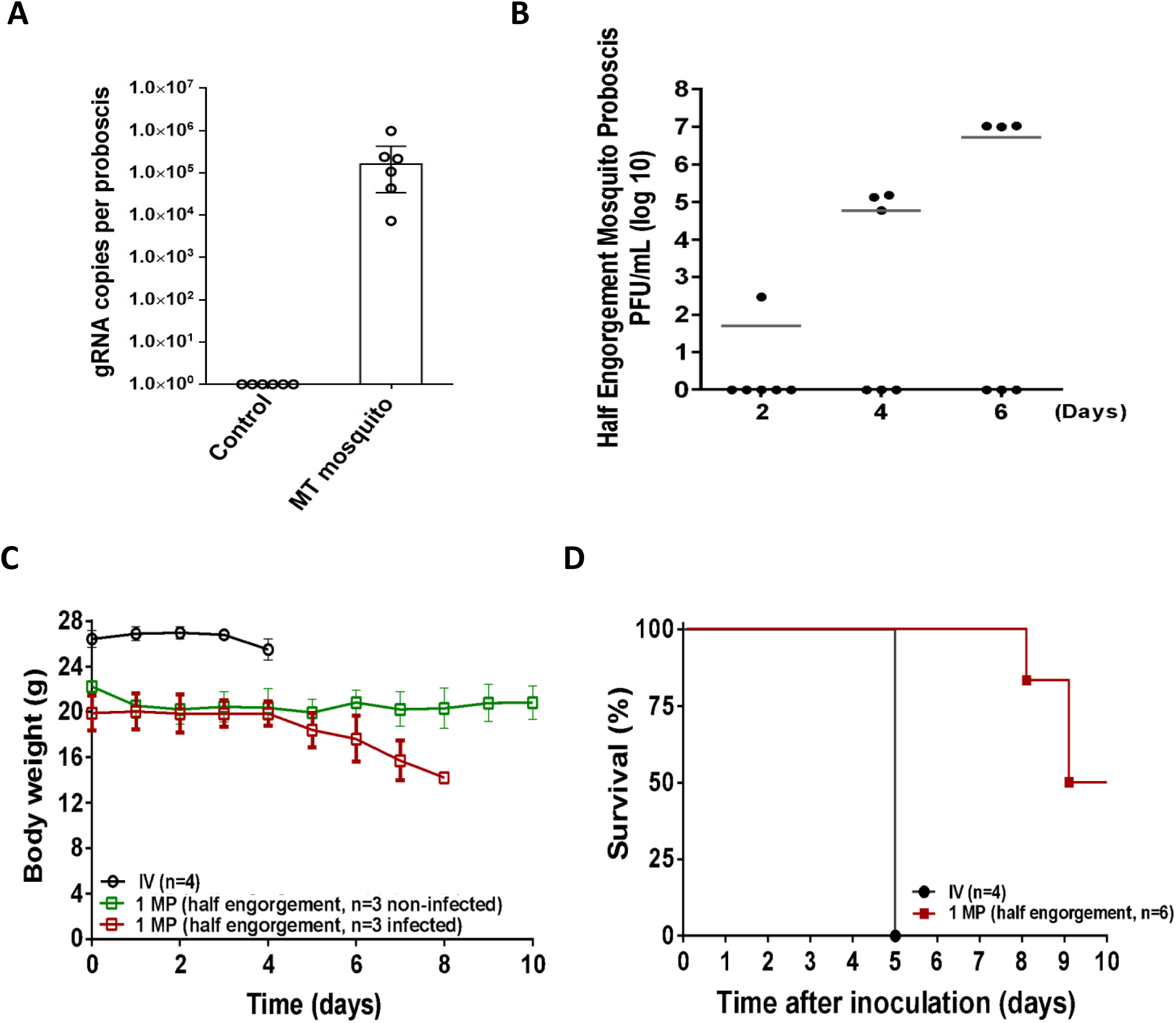
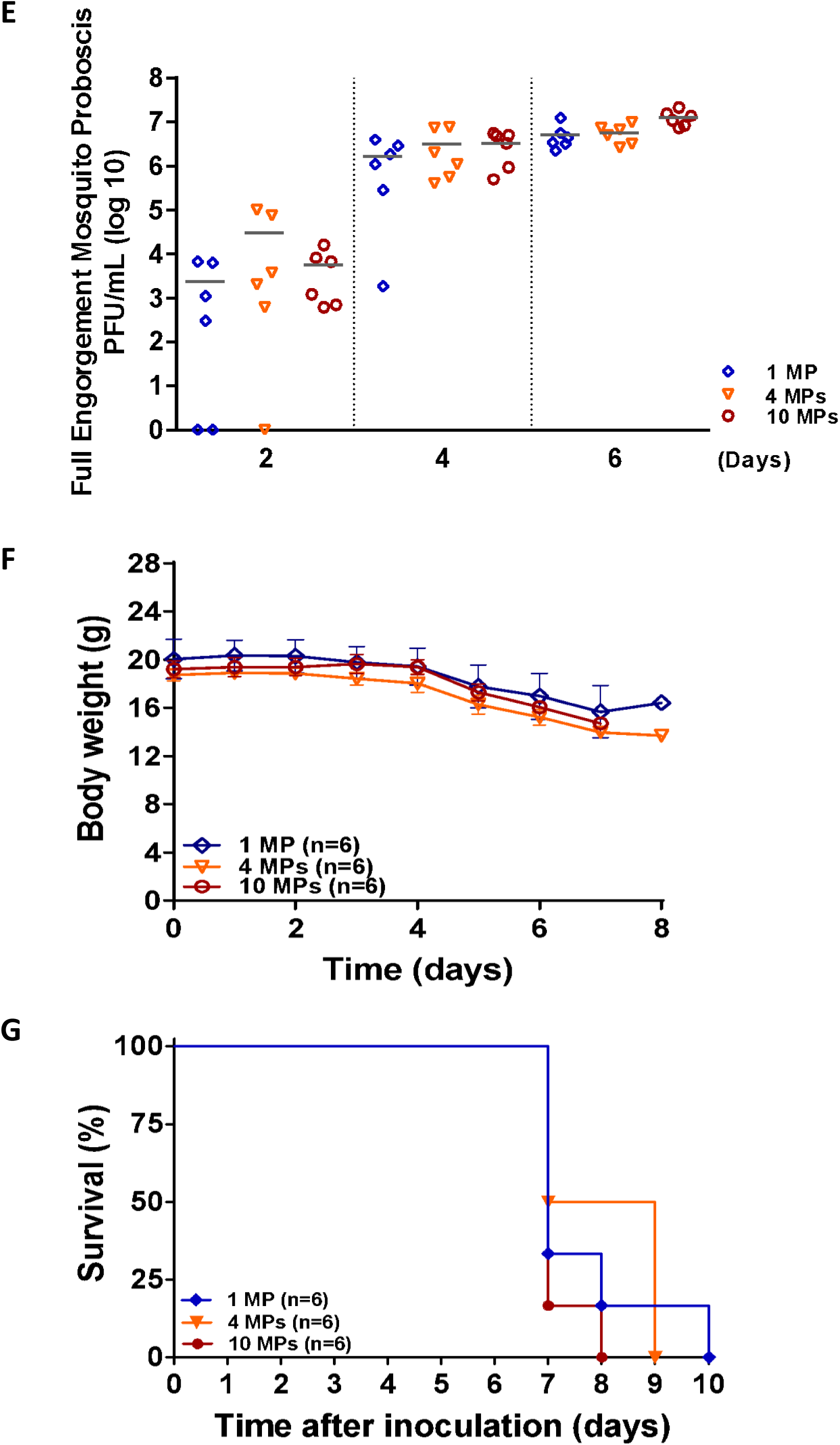
Residual DENV in proboscis of A. aegypti mediates dengue spread via mechanical transmission. (A) Copies of dengue genomic RNA in the proboscis of a mosquito after biting DENV-infected or non-infected mice. Mosquito proboscises (n = 10) were collected and examined for each experiment/group. Six independent experiments were performed. (B) Virus titer of DENV-2 in mice serum, (C) body weight, and (D) Kaplan-Meier survival curves of male mice intravenously injected with DENV (n = 4 mice) or the extracts from the indicated number of half engorged mosquito proboscises (MPs) from DENV blood-exposed A. aegypti mosquitoes (n = 6 mice per group for proboscis extract injection). (E) Virus titer of DENV-2 in mice serum, (F) body weight, and (G) Kaplan-Meier survival curves of male mice intravenously injected with the extracts from the indicated number of fully engorged mosquito proboscises from DENV blood-exposed A. aegypti mosquitoes (n = 6 mice per group for injection of proboscis extracts). The number of mice used in each group per individual experiment is highlighted.

### Mechanical transmission contributes to the speed and intensity of DENV outbreaks in mathematical models

As the proportion of DENV infections resulting from MT in human populations is unknown, we wanted to examine the potential importance of this route to determine how it may affect the course of outbreak spread. We developed a susceptible-exposed-infectious-removed (SEIR) mathematical model of single-strain dengue transmission^24^ incorporating a MT component^25^ (Supplementary Figure 3) to study these population-level dengue infection dynamics. To achieve this, we modified the vector transmission model of Esteva and Vargas^25^ to enable us to determine more accurately the relative contributions of MT and biological transmission following viral amplification in the mosquito midgut.

With this model, we examined the impact of MT at different transmission settings by exploring a variety of parameter values that led to a wide range of estimates for the total proportion of the human population infected (Supplementary Table 1; Supplementary Table 2; see Methods section for selection of parameter ranges). Supplementary Figure 4 shows the impact of mechanical transmission on population-level dengue infection dynamics under different parameter values. In particular, the total number of people infected increased by around 2.5%, with approximately 8.3% of these resulting directly from MT, when we set the average number of adult female mosquitoes per person [*k*] = 2, mosquito biting rate [*b*] = 0.5, MT probability [*p_M_*] = 0.1, number of bites [*n_M_*] = 1. When the MT probability [*p_M_*] increases from 0.1 to 1, the total number people infected increased by around 12%, with approximately 60% of these resulting directly from MT (Supplementary Table 2; Supplementary Figure 4).

Modifying the biting rate *b* to be randomly generated from 0.5 to 2 and setting MT probability *p_M_* = 0.1, 0.2, 0.4, 0.6, 0.8, 1 (Supplementary Table 2), we explored a wide range of estimates for the total proportion of the human population infected (Figures 5A-G). In this case, with *p_M_* = 0.1, almost 11% of infected individuals were infected via MT (Figures 5A, G). For a higher MT probability, *p_M_* = 0.8, the total number of infected individuals increased by 15% compared to the total number of infected individuals for *p_M_* = 0.1, 69% of whom were infected directly from MT (Figures 5E, G). For the highest MT probability, *p_M_* = 1 (the experimental value), the total number of infected individuals increased by 16.5% compared to the total number of infected individuals for *p_M_* = 0.1, 79% of whom were infected directly from MT (Figure 5F). The change in the time of peak incidence also varied considerably, with *p_M_* = 0.1 resulting in a 11 day forward shift as compared to a model without mechanical transmission (Figure 5G).

**Figure 5.**
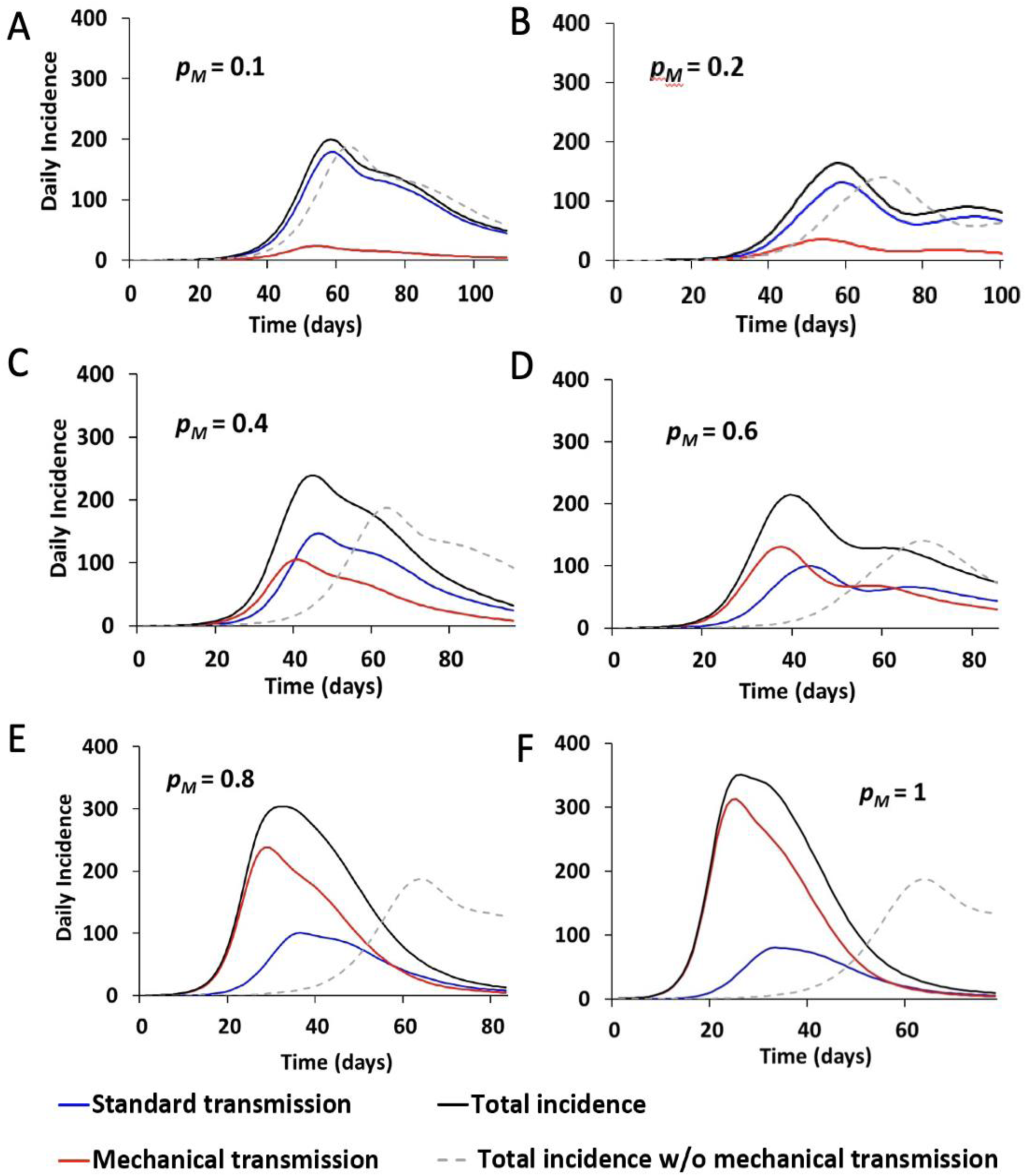
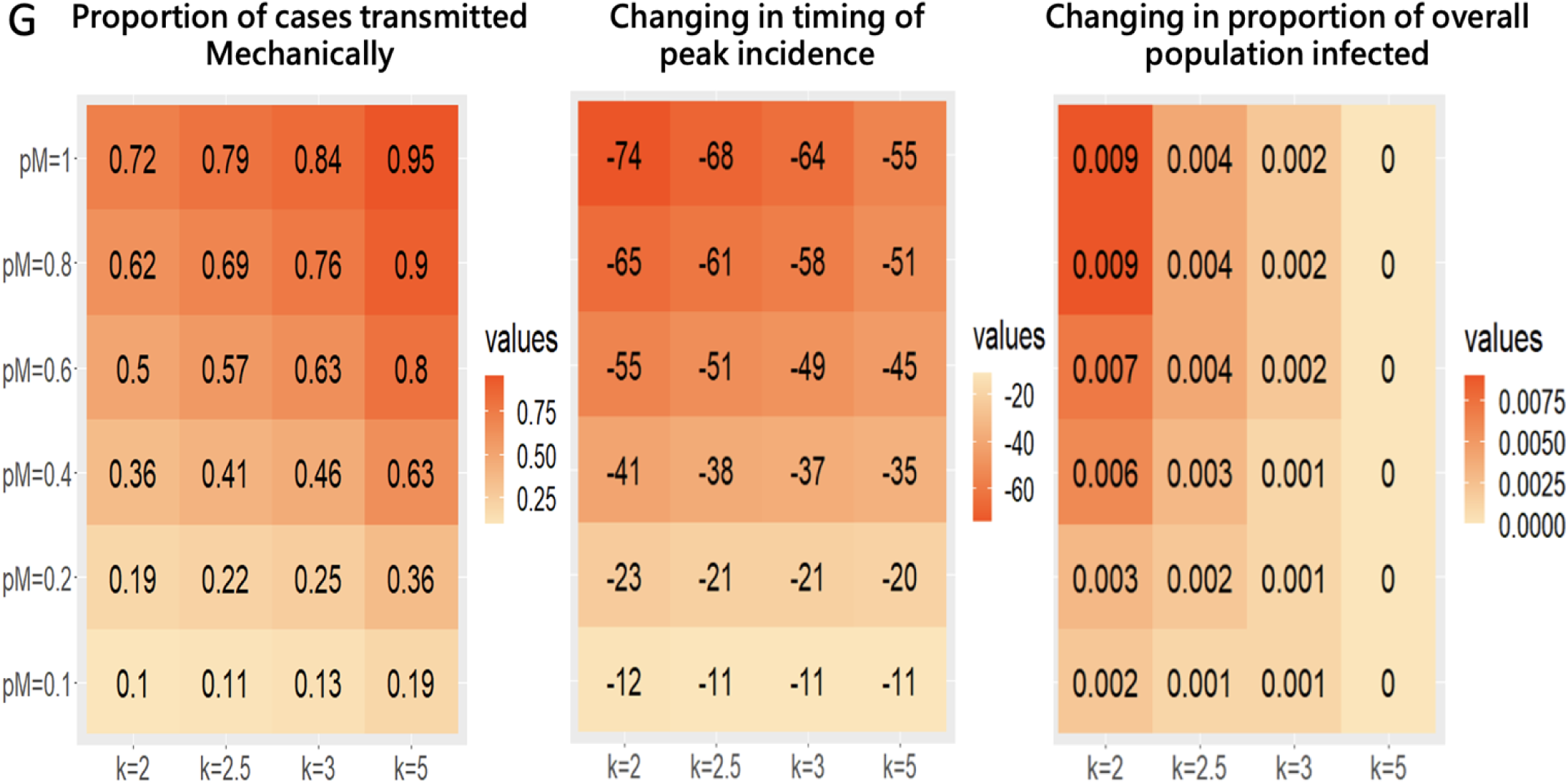
Mathematical model of dengue dynamics incorporating mechanical transmission. (A, F) Simulated daily incidence over the course of a dengue epidemic beginning with a single infected individual in an immunologically naïve population of 10,000 individuals (model in Equations 1–12 in Figure S3 and parameters in Supplementary Table 1). The average duration of MT is one hour, during which time a mosquito is expected to bite one to four other person(s), nM = 1∼4, with a probability of mechanical transmission per bite, pM, of (A) 10%, (B) 20%, (C) 40%, (D) 60%, (E) 80%, or (F) 100%. The mosquito biting rate, b, was set to be a randomly generated number from 0.5 to 2, and the average number of adult female mosquitoes per person, k, was set to 2.5. The total daily incidence (black line) is partitioned into incidence due to standard transmission following viral amplification in the vector (blue line) and incidence due to MT (red line). For comparison, the incidence when MT is not allowed (pM = 0) is shown by the gray dashed line. (G) The impact of MT on dengue infection dynamics is summarized by the proportion of cases caused by MT, the change in the timing of peak incidence, and the increase in the proportion of infected individuals after incorporating MT into the model under a range of pM and k values.

Depending on the parameters used, this change in peak incidence could range from a minimum of 11 days to 74 days earlier than for models where transmission follows only viral amplification (Figure 5G; Supplementary Figure 4). The magnitude of this peak appeared increased for models containing MT; in the outbreaks modeled in Figure 5, daily incidence can peak at around 15% greater than for the model without MT.

The total number of infected individuals also increased by between 0-14.4%, and 8.3-99.7% of dengue cases were estimated to be acquired via MT over the course of a simulated epidemic (Figure 5; Supplementary Figure 4). Thus, for a wide range of parameter values, MT is expected to influence not only the timing, but also the severity, of DENV outbreaks.

We then explored how the proportion of cases acquired through MT over the course of the epidemic changes with the number of bites during the MT period, *n_M_*, and the probability of mosquito-to-human transmission during MT, *p_M_* (Supplementary Figure 4). Even for a low MT probability, the proportion of cases transmitted by this route was non-negligible; for *p_M_* = 0.1 with *b* = 0.5, the proportion of MT cases was predicted to be 8.3–14.4% for *n_M_* = 1 and 15.9–27.4% for *n_M_* = 2. For the highest MT probability of 100% per bite, the proportion of cases resulting from MT rises to 60.1–86.7% for *n_M_* = 1 and 86.1–98.6% for *n_M_* = 2.

Taken in conjunction with the results of our DENV infection experiments, these modeling results indicate that MT of DENV is possible by *Ae. aegypti* that engage in multiple biting events. While the extent of MT in human populations is unknown, it is conceivable that it makes a non-negligible contribution to DENV transmission dynamics, especially in densely populated regions.

## Discussion

In this study, we investigated the transmission routes of dengue virus in *Ae. Aegypti* mosquitoes. Previous studies have shown that these mosquitoes can transmit DENV biologically, but the possibility of mechanical transmission has not been fully explored. Our findings show that *Ae. aegypti*-mediated DENV infection can be transmitted both biologically and mechanically. Our study used mice-based transmission assays to show that around 25% of mice can become infected with DENV from mosquitoes that had previously bitten infectious mice, similar to other bloodborne viruses like hepatitis B virus (HBV)^26, 27^. Mathematical modeling further suggests that MT may account for a significant percentage of overall transmission and shift the timing of peak disease incidence forward.

Our analyses suggest that targeting potential DENV-relevant receptors in mosquitoes^28^ may offer a control option for reducing the risk of dengue transmission. Several putative receptors or attachment factors in *A. aegypti* or mosquito cells, including prohibitin, protein R67, R80, and glycoprotein family members such as gp40 and gp45, have been associated with DENV recognition. Further research is needed to determine the minimum viral loads required for MT to occur and to understand the mechanisms involved in the interactions between DENV and the mosquito proboscis.

Prior studies from decades ago have demonstrated the potential for MT to play a role in transmitting *Aedes*-borne viruses such as Chikungunya. More recent studies have identified viral CHIKV and DENV on the proboscises of female *Ae. aegypti* immediately after uptake of an infectious blood meal, though this report found no evidence of MT playing a role in CHIKV transmission (in direct contradiction to earlier studies, and potentially due to the strain of mouse used)^29, 30^. Our study strongly supports MT as playing a role in dengue transmission via DENV-contaminated proboscises; in particular, MT could be important as a potential driver of rapid dengue outbreaks.

Recent work demonstrating the increased attractiveness of mice infected with flaviviruses to host seeking *Aedes* females may also be relevant from a MT perspective, as this could increase the likelihood of female mosquitoes first biting those infected before completing their blood meal (and potentially transmitting disease) by biting non-infected individuals^31^. This suggests individuals within the same household are potentially not equally likely to be bitten, supporting an increased speed of disease transmission. With our current experimental setup, we were unable to exclude the possibility of the influence of time intervals between bites by infected mosquitoes on MT efficiency, resulting in potential changes in load or longevity of residual DENV.

There are several limitations to this study that need to be addressed in future experiments. For example, the use of immunocompromised mice limits the generalization of our results to the human population. In addition, the influence of time intervals between bites by infected mosquitoes on MT efficiency, and the residual load or longevity of DENV, remains unclear. Despite these limitations, our results have important implications for the control of dengue outbreaks and highlight the need for further research into the role of MT in dengue transmission.

## Materials and Methods

### Virus and cells

The clinical isolate 2015 DENV-2 (TW2015)^12^ was propagated in *Aedes aegypti* and Vero cells, and the viral titer was determined via plaque-formation assay using BHK- 21 C15 cells. Vero cells were grown at 37°C, 5% CO2 in 1× Dulbecco’s modified Eagle’s medium (DMEM) supplemented with 5% fetal bovine serum (FBS), 1× MEM nonessential amino acid solution, and 1× antibiotic-antimycotic solution. BHK-21 C15 cells were grown in 1× DMEM supplemented with 2% FBS and 1× antibiotic-antimycotic solution at 37°C, 5% CO2. All reagents were purchased from Thermo Fisher Scientific.

### Plaque-formation assay

The viral titer of mouse serum stored at -80°C was determined by a previously described plaque-formation assay^32^. In brief, virus-containing serum samples from mice were serially diluted in serum-free DMEM and added to a BHK-21 C15 cell monolayer for virus adsorption at 37°C for 2 hours. After the diluted samples were removed, the BHK-21 C15 cells were overlaid with DMEM containing 1% methylcellulose (4000 cps, Sigma-Aldrich), 5% FBS, 2 mM glutamine, 1 mM sodium pyruvate, 2.5 mM HEPES, and 1× antibiotic-antimycotic solution and cultured for 6 days at 37°C. The overlaid medium was then gently removed, and the cells were stained with Rapid Gram Stain solution (Tonyar Biotech) for 2 hours to stain plaques before being washed with water. Plaques were counted, with viral titers expressed as plaque-forming units (PFU)/mL.

### Mosquito husbandry

*Ae. aegypti* (Higgs strain) mosquito eggs were hatched in deionized water under deoxygenated conditions. Hatched larvae were fed powdered yeast and goose liver (NTN Fishing Bait). Newly emerged mosquitoes were collected in rearing cages and fed a 10% sucrose solution for maintenance at 28°C in 70% relative humidity with a 12-hour light/dark cycle. Five-day-old female mosquitoes were starved for 16 hours prior to mechanical DENV transmission experiments.

### Mouse husbandry

All animal experiments were conducted in compliance with the Guidelines for the Care and Use of Laboratory Animals published by the National Research Council, Taiwan (1996). The animal protocol was approved by the Institutional Animal Care and Use Committee of the National Health Research Institutes (NHRI-IACUC-107054-A). AGB6 mice were provided and genotyped by Dr. Guann-Yi Yu’s lab (NHRI, Taiwan). All mice were bred and maintained at the ABSL-2 animal feeding room of the Laboratory Animal Center of the NHRI at 22 ± 2°C in 55 ± 10% relative humidity with a 12-hour light/dark cycle. The Center received full accreditation from the Association for Assessment and Accreditation of Laboratory Animal Care (AAALAC) International in 2015. All food, water, aspen chips, and cages were sterilized before use. Mouse health monitoring was performed every 3, 6, or 12 months by Serology, Microbiology and Parasites, using the standard operating protocol of the NHRI Animal Center (https://lac.nhri.org.tw/08-2/).

### Infection of AGB6 mice with DENV

Eight- to ten-week-old, 18–24 g female and/or 22–25 g male AGB6 mice were challenged intravenously with either 1000 PFU of DENV^33^ in 100 μL of serum-free DMEM or with 1× PBS (control) three days before the start of all experiments. For DENV infection via injection of mosquito proboscis extracts, female *Ae. aegypti* were first allowed to feed from DENV-infected mice until either half or fully engorged. Mosquito proboscises were collected and dissected in groups of one, four, or ten; these were then ground in 100 µL of 1× PBS prior to inoculating naïve AGB6 mice via intravenous injection.

### Infection of AGB6 mice with DENV via mechanical transmission

AGB6 mice were intraperitoneally injected with the anesthetics Rompun(16 mg/kg, Bayer Animal Health) and Ketalar (100 mg/kg, Pfizer). Anesthetized mice were individually placed on top of the mesh covering a mosquito cage and exposed to three to five mosquitoes that had been starved for 16 hours. These mosquitoes were allowed to feed from DENV-infected AGB6 mice until they were half-engorged, as confirmed via visual observation. The mice were then immediately removed and naïve AGB6 mice were placed on the top of the cage. The mosquitoes were allowed to resume feeding until fully engorged. The number of successful mosquito bites was determined by counting the number of blood-engorged mosquitoes. Mouse serum was collected by retro-orbital bleeding on days 0, 2, 4, and 6 after mosquito biting, and body weight and survival were monitored for 10 days.

### Quantification of DENV genomic RNA

Four days after biting non-infected/DENV-infected mice, the mosquitoes were dissected. The proboscis tissues were collected, and RNA was extracted using a standard Trizol-based protocol. Extracted RNA was reverse transcribed to cDNA using a SuperScript III Reverse Transcriptase Kit (Thermo Fisher Scientific) prior to real-time quantitative PCR (RT-qPCR) analysis. RNA fragments encompassing the qPCR target were produced and used to generate an absolute standard curve as described previously^34^. DENV genomic RNA was quantified using a RT-qPCR detection system (ABI) with a thermal profile of 95°C for 3 min, 95°C for 2 sec, and 40 cycles of 60°C for 20 sec. The following primers were used for RT-qPCR analysis in this study:

DENV2 forward primer: 5’-TCG CTG CCC AAC ACA AG-3’

DENV2 reverse primer: 5’-CAT GTT CTT TTT GCA TGT GAA C-3’.

### Mathematical modeling

We created a mathematical model of DENV transmission, including both a susceptible-exposed-infectious-removed (SEIR) model for human transmission and a susceptible- mechanical transmitting-exposed-infectious (SMEI) model for mosquito transmission, where “M” represents the time period during which females could potentially mechanically transmit DENV following uptake of an infectious blood meal (Supplementary Figure 3).

We assume that humans enter the population as susceptible, *S_H_*, and describe this population by

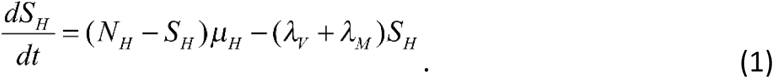

Here, *SH* is the number of susceptible humans in the total population, *μH* is the birth rate (set as equal to the death rate), and *NH* is the total human population size. The force of infection (the probability of an infection event per person per unit time) is split into two components, one part resulting from standard disease transmission following viral amplification in the mosquito, *λ_V_*, and a second part resulting from MT, *λ_M_*. We estimate the force of infection after viral amplification in the mosquito by

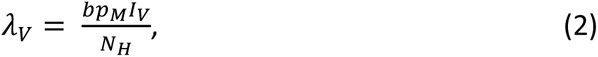

where *I_V_* is the number of DENV-infectious mosquitoes, *b* is the biting rate of mosquitoes, and *p_M_* is the probability of mosquito-to-human transmission occurring when an infectious mosquito bites a susceptible human. Similarly, we estimate the force of infection resulting from MT by

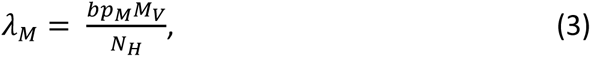

where *M_V_* is the number of mosquitoes that have taken a blood meal and have the potential to transmit DENV mechanically, *b_M_* is the biting rate of these mosquitoes, and *p_M_* is the probability of mosquito-to-human transmission when a mosquito in this category bites a susceptible human.

Once susceptible humans have become infected, they progress to the latently-infected group (those that are infected but not yet infectious), *E_H_*, which is described by

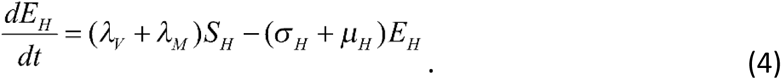

Here, *EH* is the number of latently infected humans in the total population, and *σH* is the progression rate by which an individual changes from being latently infected to being infectious. The population of infectious individuals, *IH*, is described by

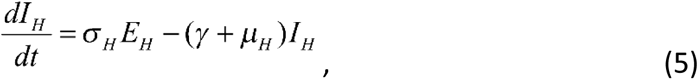

where *γ* represents the rate at which an infectious individual recovers. These recovered individuals cannot be infected again, and the recovered population group is described by

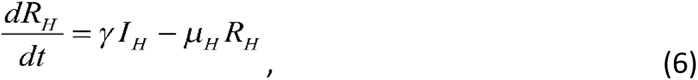

where *R_H_* is the population of recovered humans. This model ignores other DENV- relevant factors, such as antibody-dependent enhancement and disease-induced mortality.

Mosquito population dynamics are modelled by a related set of equations. We assume that all female mosquitoes are born susceptible and that this susceptible population, *S_V_*, can be described by

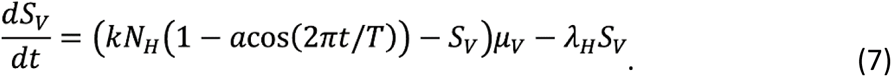

Here, *S_V_* is the susceptible vector population, *k* is the average number of adult female mosquitoes per human, *μ_V_* is the female mosquito mortality rate, *T* is the seasonal period (in this case, 1 year), and *a* is the degree of seasonality influencing the female mosquito population size (*a* = 0 in the absence of seasonality, and *a* = 1 if the peak mosquito emergence rate is twice the mean and the minimum emergence rate is 0). The force of infection, *λ_H_*, governing infection of mosquitoes by humans (when susceptible females bite infectious humans) is described by

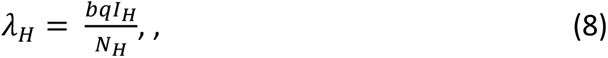

where *b* is the mosquito biting rate, and *q* is the probability of human-to-mosquito transmission. Before becoming latently infected, female mosquitoes pass through a MT stage, *M_V_*, described by

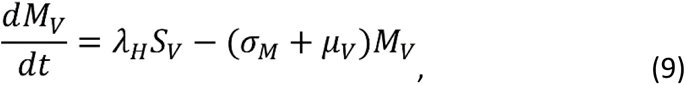

where *M_V_* is the number of female mosquitoes in the MT group, and *σ_M_* is the inverse of the duration of this phase. Following a latency period of duration *τ_V_*, mosquitoes enter an infectious stage (via standard transmission rather than MT), *I_V_*, described by

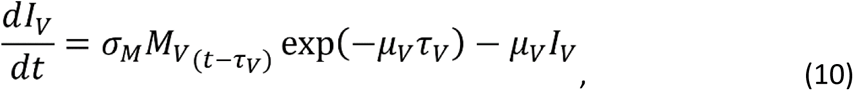

where *I_V_* is the number of infectious females. We use a delay formulation to model the latently infected phase because the latent period is long relative to the lifetime of a mosquito (10 day latent period versus 14 day mean adult female lifespan ^35^), hence an exponentially distributed time spent in the latently infected stage overestimates the number of vectors that are infectious following viral incubation within the mosquito. We assume that female mosquitoes cannot recover after infection, and thus remain infectious until death. The number of latently infected mosquitoes, *E_V_*, is described by

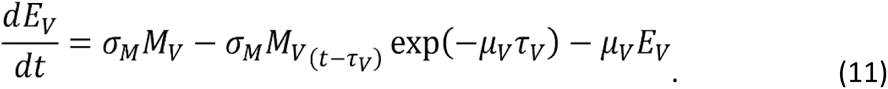

Here, we have neglected vertical transmission of dengue fever among vectors (from parent to offspring).

Parameter estimates were sourced from previously published literature wherever possible (Supplementary Table 1), while MT parameters were based on results generated by this study. Select parameters were then explored in more depth via sensitivity analyses. The duration of MT, 1/*σ_M_*, was estimated at 1 hour. The number of humans bitten during the MT period, *n_M_*, was also estimated; we assumed that a female mosquito would on average bite one human during the MT period immediately following her initial bite of an infectious human. We thus estimated the MT period biting rate to be

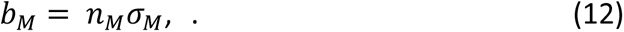

The probability of mosquito-to-human transmission for bites received by humans during the MT period was based on the experiments conducted on immunocompromised mice. Here, 16% and 40% of mice were infected after receiving two and four infective mosquito bites, respectively. We assumed that each mosquito bite has an equal and independent probability of infection. This allowed for the probability of transmission after *k* bites, *p_k_*, to be described as

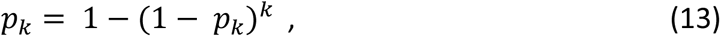

where *p_M_* is the probability of transmission for each bite. Hence, *p_M_* can be calculated as

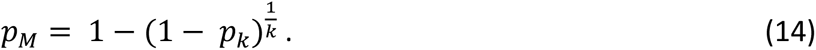

Equation 14 yields a transmission probability of 0.12 per bite for the case of two infective bites, and of 0.083 per bite for the case of four infective bites, the mean of which is 0.10 per bite. We treated this transmission probability as an upper limit for our analyses, and also considered a range of lower values (0.05, 0.025, 0.0125) in simulations to investigate potential differences between animal experiments and human transmission.

The initial parameters for each simulation were set as *IH* = 10, *IV* = 100, *NH* = 10000, and 0 for all other states. Furthermore, we consider the following parameters randomly generated from a range.

1. The average latent period in human host 1/σ*_H_* is a randomly generated number from 3 to 5 days.
2. The average latent period in mosquito host *τ_v =_* 1/σ*_v_* is a randomly generated number from 7 to 11 days.
3. The average infectious period in human host 1/γ is a randomly generated number from 3 to 6 days.
4. The mosquito biting rate *b* is a randomly generated number from 0.5 to 2.
5. The number of bites during mechanical transmission period *n_M_* is randomly chosen from the set {1, 2, 3, 4} with equal probability. The code itself is available in a GitHub repository (https://github.com/roach231428/mechanical_transmission).

### Statistical analysis

Mosquitoes were randomly selected from cages, and mice were assigned to different groups according to average weight. A significance level of *p* < 0.05 was used throughout (\**p* < 0.05; \*\**p* < 0.01; \*\*\**p* < 0.001; \*\*\*\**p* < 0.0001). Mosquito and mouse serum titers were tested using non-parametric Mann–Whitney tests. Mouse body weights were tested using parametric unpaired t-tests. A log-rank (Mantel-Cox) test was used to compare survival distributions. Four biological replicates were conducted for all experiments. Statistical analysis was conducted using the GraphPad Prism 6 statistical software.

## Acknowledgments

The authors would like to thank the NHRI Laboratory Animal Center for mouse rearing and health monitoring, and Anthony A. James for his advice on the manuscript. The authors thank Ms. Fang-Jing Lee and Mr. Yi-Kai Chen of the Institute of Population Health Sciences, the National Health Research Institutes, Taiwan, for their help with data management.

## Data sharing plans

All data collected, as well as data analysis scripts, will be made available via Dryad and GitHub upon publication.

## Funding information

This work was supported by grants to CHC and GYY from the NHRI (04D2- MMMOST02) and the Ministry of Science Technology (MOST104-2321-B-400-016).

## Notes

### Competing Interest Statement

The authors have declared no competing interest.

