## Supplementary figures and tables for "Mechanical transmission of Dengue Virus by *Aedes aegypti* may influence disease transmission dynamics during outbreaks"

### Supplementary figure 1

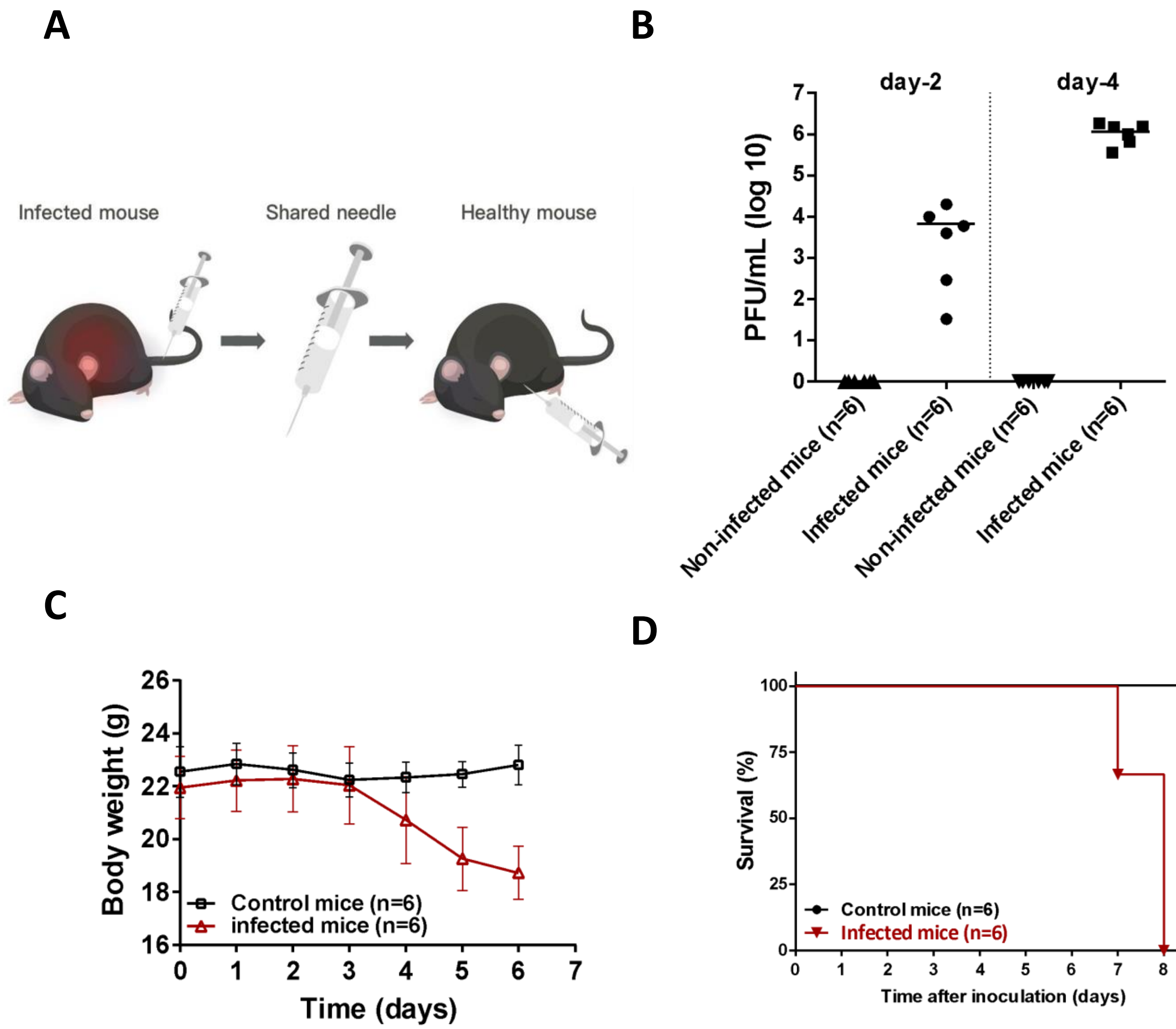

**Figure S1. Dengue transmitted by a way of needle sharing causes viremia and mortality in mice.**

(A) Schematic representation of DENV transmission via a shared needle sticking from dengue-infected mice. (B-D) A naïve AGB6 mice intravenously injected with DENV (1000 PFU) at 6 dpi is used for creating a DENV-contaminated needle with sticking tail vein of the DENV-infected AGB6 mice. Virus titer of DENV in serum (B), body weight (C) and survival (D) in male AGB6 which were intraperitoneally stuck with DENV-contaminated needle (n=6 mice, infected) or with  sterile needle (n=6, control) for totally four times.

#### Supplementary figure 2

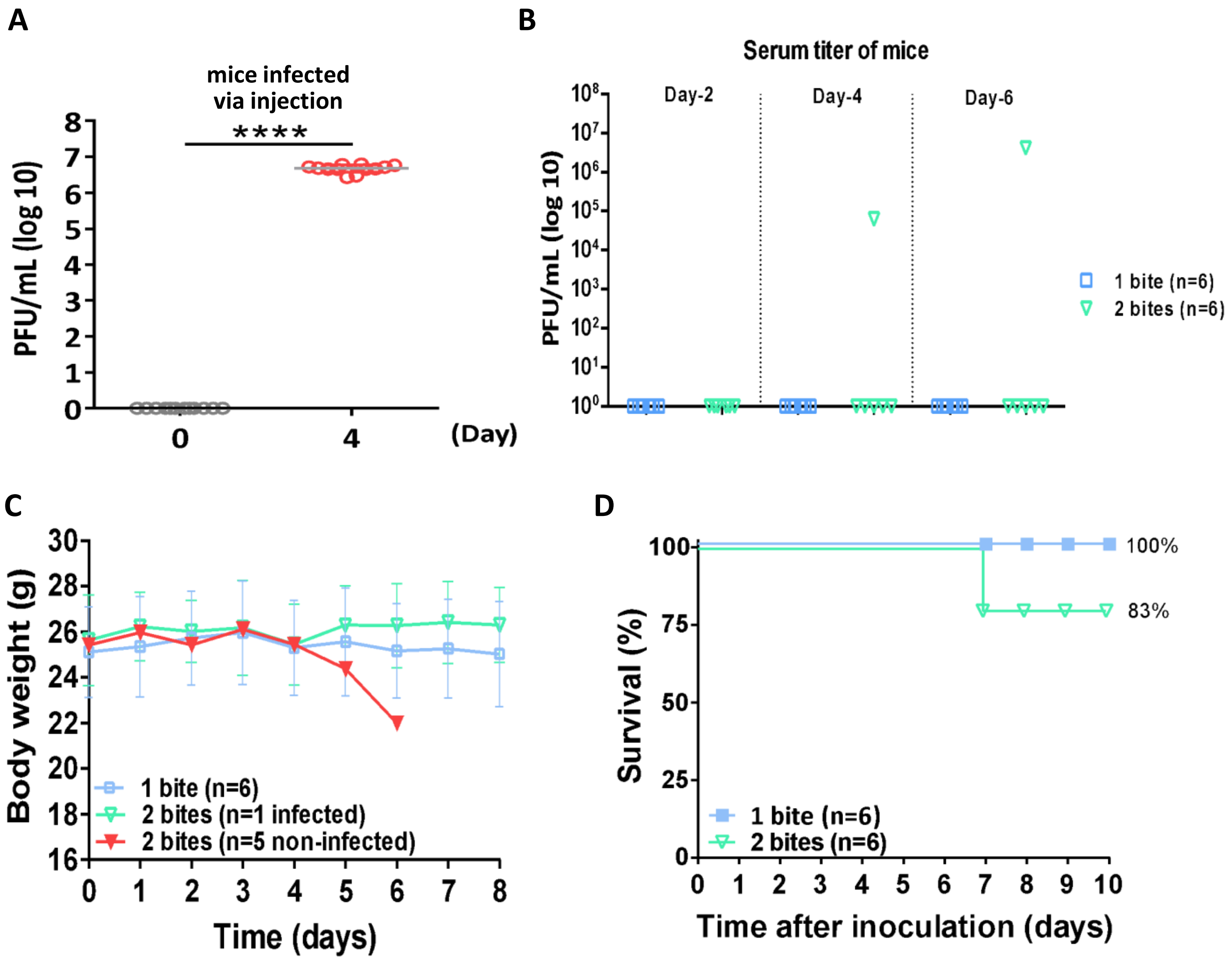

**Figure S2. Related to Figure 1. A pre-test of effects of AGB6 mice caused by dengue mechanical transmission via *Aedes* mosquitoes.**

(A) DENV titer in the serum of female and male mice was examined by plaque-formation assay at four days after intravenous infection with DENV. Circles represent means of virus titer from DENV-inoculated AGB6 mice (n=12). (B-D) Serum DENV (B), body weight (C), and survival (D) were examined in female (n=6 for each group) and male (n=6 for each group) mice potentially infected with DENV via mechanical transmission by one or two bites of DENV blood-sucked *A. aegypti.*

**Supplementary figure 3**

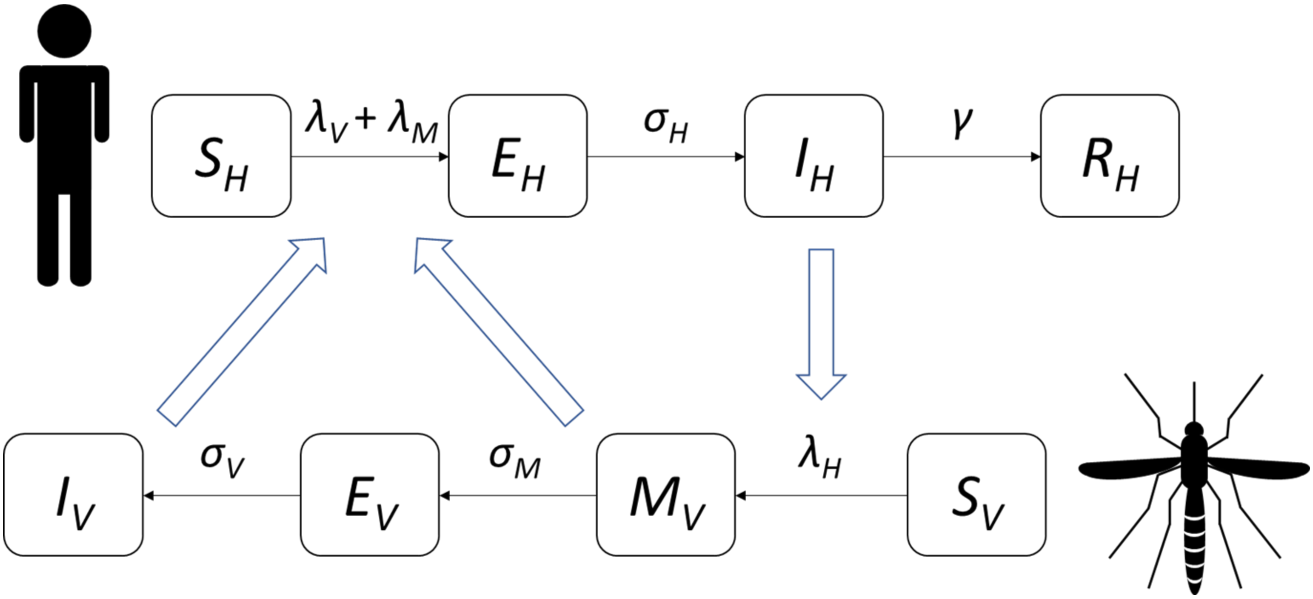

**Figure S3. Mathematical model of dengue transmission dynamics including a mechanical transmission component.**

Once susceptible humans, $S_H$, have become exposed to the virus, they become latently-infected with a force of infection comprised of a mechanical transmission component, $\lambda_M$, and a viral amplification component, $\lambda_V$. These latently-infected humans, $E_H$, then become infectious after an average latency period of $1/\ \sigma_H$. Infectious humans, $I_H$, can recover after an average infectious period of $1/\gamma$. Upon recovery, recovered humans, $R_H$, are assumed to be immune to further infection events. Similarly, susceptible vectors, $S_V$, become infected dependent on the force of infection due to infectious humans, $\lambda_H$. They can then briefly transmit disease whilst in a mechanical transmission stage, $M_V$, whose duration is given by $1/\sigma_M$. These females enter a latently-infected category, $E_V$, where they remain for a duration equal to $1/\sigma_V$. After these females become infectious to humans, $I_V$, they remain so for the duration of their lifespan.

### Supplementary figure 4

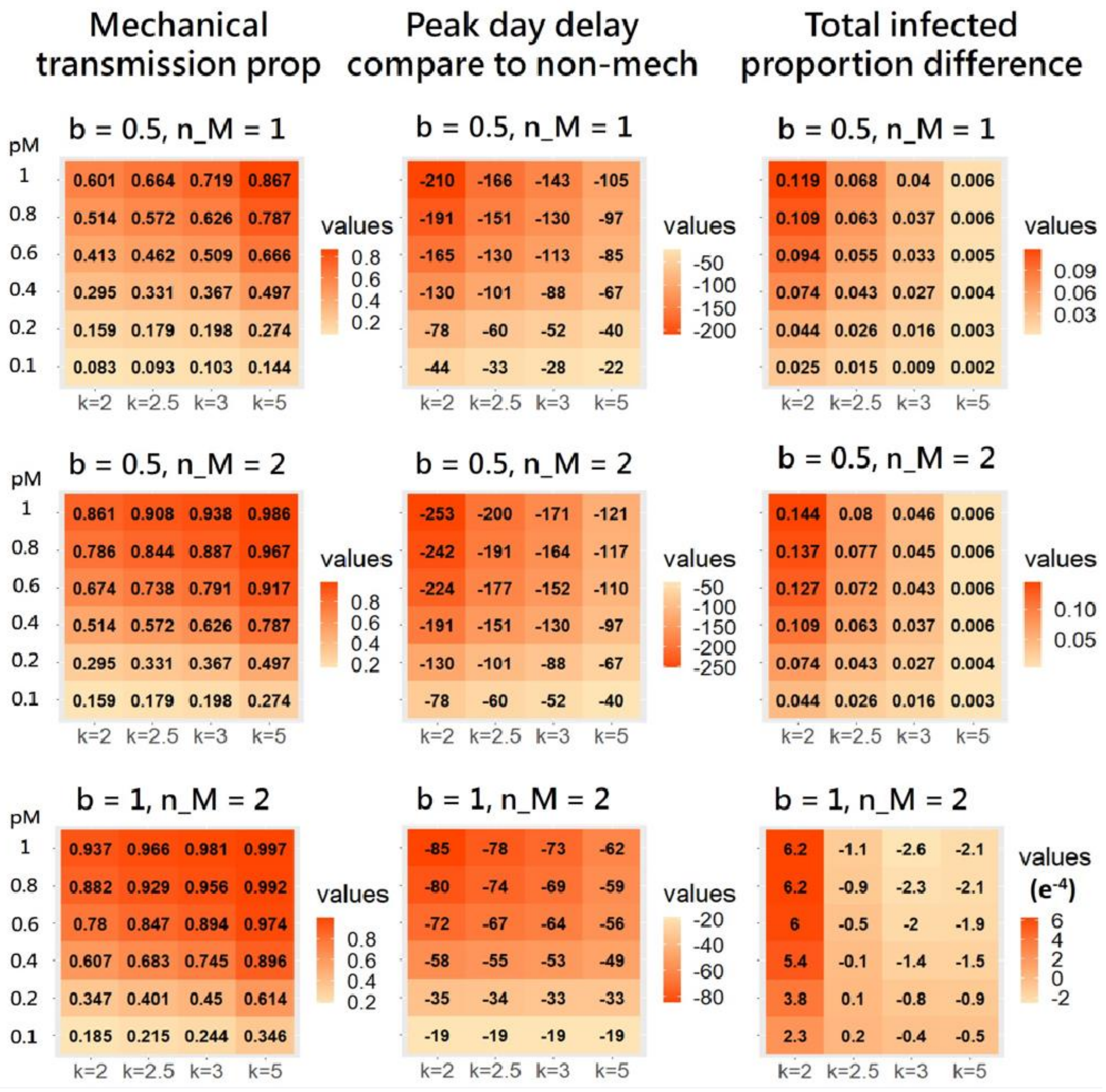

**Figure S4. The impact of mechanical transmission on population-level dengue infection dynamics under different parameter values.**

Impact of altering key parameter values on dengue infection dynamics summarized by (left) the proportion of cases caused by MT, (centre) the change in the timing of peak incidence, and (right) the increase in the proportion of infected individuals after incorporating MT into the model. k = number of adult female mosquitoes per human; pM = MT transmission probability; b biting rate; nM = number of bites during MT period.

**Table S1. Table of parameter estimates used in the dengue transmission model.**

Ranges for variable parameters are shown in parentheses.

| Parameter | Description | Value: | Reference |
|---|---|---|---|
| $1/\mu_H$ | Average adult human lifetime | 70 years | |
| $N_H$ | Human population size | 10,000 | |
| $k$ | Average number of adult female mosquitoes per person | 2, 2.5, 3, 5 | (1), (2) |
| $1/\mu_V$ | Average adult female mosquito lifetime | 14 days | (3), (4), (5) |
| $1/\sigma_H$ | Average latent period in human host | randomly generated from 3 to 5 days | (6), (7) |
| $\tau_V = 1/\sigma_V$ | Average latent period in mosquito host | randomly generated from 7 to 11 days | (6), (8) |
| $1/\gamma$ | Average infectious period in human host | randomly generated from 3 to 6 days | (9), (10) |
| $b$ | Mosquito biting rate | randomly generated from 0.5 to 2 times per day | (4), (11) |
| $p$ | Mosquito-to-human transmission probability (following parasite incubation) | 0.38 | (12) |
| $q$ | Human-to-mosquito transmission probability | 0.38 | (12) |
| $a$ | Degree of seasonality in mosquito population size | 0 | |
| $T$ | Period of seasonality | 1 year | |
| $1/\sigma_M$ | Average duration of mechanical transmission | 1 hour | |
| $n_M$ | Number of bites during mechanical transmission period | 1, 2 | |
| $p_M$ | Mosquito-to-human transmission probability during mechanical transmission | 0.1, 0.2, 0.4, 0.6, 0.8, 1 | |

**Table S2. Changes in expected proportion of human population infected for a range of model parameters.**

| ($b$, $n_M$) $k$ | (0.3, 1) | (0.5, 1) | (0.3, 2) | (0.5, 2) |
|---|---|---|---|---|
| 2 | 11.27%* | 91.36% | 11.27% | 91.36% |
| 2.5 | 43.67% | 95.54% | 43.67% | 91.54% |
| 3 | 62.05% | 97.56% | 62.05% | 97.56% |
| 5 | 89.36% | 99.65% | 89.36% | 99.65% |

*Similar to the 2015 outbreak in Kaohsiung.
